# Global maps of transcription factor properties reveal threshold-based formation of DNA-bound and mobile clusters

**DOI:** 10.1101/2025.04.24.650477

**Authors:** Sadia Siddika Dima, Meghan A. Turner, Hernan G. Garcia, Gregory T. Reeves

**Author notes:** To whom correspondence should be addressed: 100 Spence Street, Texas A&M University, Chemical Engineering Dept College Station, Texas 77843 MS 3122.

## Abstract

The relationship between bulk transcription factor concentration and DNA binding has been a central question in gene regulation for decades. Recent studies propose that DNA-bound transcription factor hubs, or clusters, aid in fast and precise transcriptional interpretation. Using live imaging techniques, we quantify the concentration, binding, and mobility of the morphogen Dorsal (Dl), both in bulk and in clusters, in *Drosophila* blastoderm embryo. Our experiments encompass multiple length and time scales, allowing us to obtain a nucleus-wide view of the mechanism connecting hub formation to bulk Dl concentration. Our results show that previously unobserved, slowly-moving clusters of Dl are present, in addition to the expected populations of freely mobile and DNA-bound Dl. Furthermore, both mobile clusters and DNA-bound Dl appear only once a threshold concentration in the nucleus is surpassed, a behavior consistent with liquid-liquid phase separation. Thus, our work elucidates how bulk transcription factor concentration dictates the formation and spatiotemporal changes of different populations needed for gene regulation.

## Introduction

Cell fate determination and maintenance in metazoans depend on the accurate spatiotemporal regulation of gene expression. The binding of TFs to regulatory motifs in the DNA in eukaryotes, and thereby the target gene expression, is dependent on a plethora of factors including its concentration, binding affinity, chromatin accessibility, and protein-protein interactions (*1–4*) Therefore, bulk concentration, a parameter commonly estimated from the fluorescence intensity of fixed or live imaging, cannot serve as a direct proxy to TF activity; instead, measurements of TF-DNA binding, in relationship to the bulk concentration, are required. Recent studies also suggest TFs form DNA-bound clusters (or hubs) of elevated concentration (reviewed in (*5, 6*)), potentially mediated by intrinsically disordered regions (*7–9*). It has been proposed that these clusters represent liquid-liquid phase-separated droplets; although that suggestion remains a matter of debate (*5, 6*). In *Drosophila*, the formation of TF hubs has been found to facilitate binding to low-affinity sites by increasing the local concentration of TFs (*10–12*). These hubs aid in transcriptional burst induction (*9*) and function in a gene-specific, or enhancer-specific manner (*13*). However, the mechanisms connecting bulk transcription factor concentration, on the micron scale, to DNA binding and hub formation, on the nanometer scale, remain poorly understood.

Dorsal (Dl), a *Drosophila* homolog of mammalian NF-κB, acts as a morphogen to pattern the dorsal-ventral (DV) axis of the *Drosophila* blastoderm-stage (1-3 hr old) embryo, regulating hundreds of genes (*14–16*). Dl is one of the earliest known morphogens (*17*) and one of the best characterized sequence-specific TFs (*18*). Previous work using fluorescent imaging of live and fixed embryos indicated that the Dl gradient has a Gaussian-like shape with the highest level of Dl present on the ventral side of the embryo (*14, 19*). The level of Dl oscillates in a progressively increasing saw-tooth pattern during blastoderm nuclear cycles (ncs) 10 to 14 (*14, 20*). This temporal variation has been proposed to ensure the accurate dynamic expression patterns of its target genes (*14*). However, modeling efforts that take spatiotemporal variation of the Dl gradient as the only input fail to predict expression pattern and dynamics of all the target gene domains (reviewed in (*21, 22*)). As such, knowing the bulk nuclear intensity of Dl is not sufficient to explain patterning. Intriguingly, recent groundbreaking work has shown that the activity of many transcription factors, including Dl, is also influenced by the presence of clusters (also referred to as hubs, aggregates, droplets, or condensates), and the enrichment of Dl clusters has been observed at the site of transcription in both fixed and live embryos (*13, 23*). But how the dynamic Dl gradient dictates the formation of these populations of clusters remains unexplored.

Thus, to understand the transcriptional interpretation of the Dl gradient, knowing the dynamics of Dl bulk nuclear concentration is an important, necessary piece, but it is not sufficient by itself. In the same way, knowing the cluster formation dynamics at specific enhancers is also required for a detailed view of gene regulation (*13*), but it is again not the full picture. Measurements of different populations of Dl -- such as the freely diffusing population, the DNA-bound population, clusters of Dl -- and, in particular, how they relate to each other in dose/response maps, must be considered. Therefore, we used advanced live imaging and analysis techniques, such as raster image correlation spectroscopy (RICS) (*24–29*), a type of scanning fluorescence correlation spectroscopy (*30–32*), and single particle tracking (SPT) of super-resolution images of an endogenously-tagged Dl. These experiments, which span a wide range of length and time scales, indicate the presence of three different populations of Dl: freely moving Dl, DNA-bound Dl, and a previously unobserved population of slowly-moving clusters of Dl. Interestingly, a recent study similarly observed the formation of mobile clusters of two other blastoderm patterning factors, Bicoid (Bcd) and Capicua (Cic), and proposed these mobile clusters to be an efficient search strategy for regulatory regions (*33*). Thus, mobile cluster formation might be a common mechanism for the accurate transcriptional interpretation of morphogen gradients. Our results further suggest that the total nuclear concentration of Dl must reach a threshold of roughly 30 nM before DNA binding occurs and before the slowly-moving clusters of Dl are formed, potentially indicating an ultrasensitive process or, alternatively, a liquid-liquid phase separation (LLPS) process. Ultimately, our results suggest that the relationships of the different populations of Dl in the nucleus, including DNA-bound as well as slowly-moving clusters, must be taken into account to understand transcriptional regulation on a tissue-wide level. We suggest that the importance of these relationships is also extended to other transcription factor systems.

## Results

### The Dl nuclear concentration and binding is dynamic throughout the blastoderm stage

Studying the dose/response behavior between the bulk concentration of a TF and its DNA binding interactions, which can provide important insights into gene regulation, requires measurements at a range of concentrations. To do so, we leveraged the fact that the Dl gradient has dynamically varying levels over nc 10-14. We made use of a CRISPR-edited *dl-mNeonGreen* (*dl-mNG*) construct in which the *mNeonGreen* was inserted scarlessly in-frame just before the stop codon of the endogenous *dl* locus (referred to as the “wildtype” construct). At two copies, this scarless CRISPR edit is viable and results in similar target gene expression to wildtype. We imaged the ventral-most nuclei (Fig. 1A) from nc 10 to gastrulation in embryos collected from mothers having one copy of the Dl-mNeonGreen wildtype construct and one copy of His2Av-RFP (to mark the DNA). This line is used in the rest of our experiments unless mentioned otherwise. Our time series image acquisitions (Movie S1) were compatible with RICS analysis (see Methods), which allowed us to quantify the temporal dynamics of the total nuclear concentration and binding of Dl. We found the absolute nuclear concentration of Dl increases from one nuclear cycle (nc) to the next from ~ 60 nM in nc 10 to ~120 nM in nc 14 (blue “wt” data in Fig. 1B), consistent with the increase in Dl gradient amplitude over time observed previously (*14, 19, 20, 34*). During each nc, the nuclear concentration of Dl increases in the beginning of interphase as Dl enters the nucleus after mitosis, then drops abruptly as the nuclear envelope breaks down before the next mitosis (*14, 20*). The long duration of nc 14 compared to the other ncs allows the nuclear concentration to reach a steady state value of ~ 120 nM.

**Figure 1:**
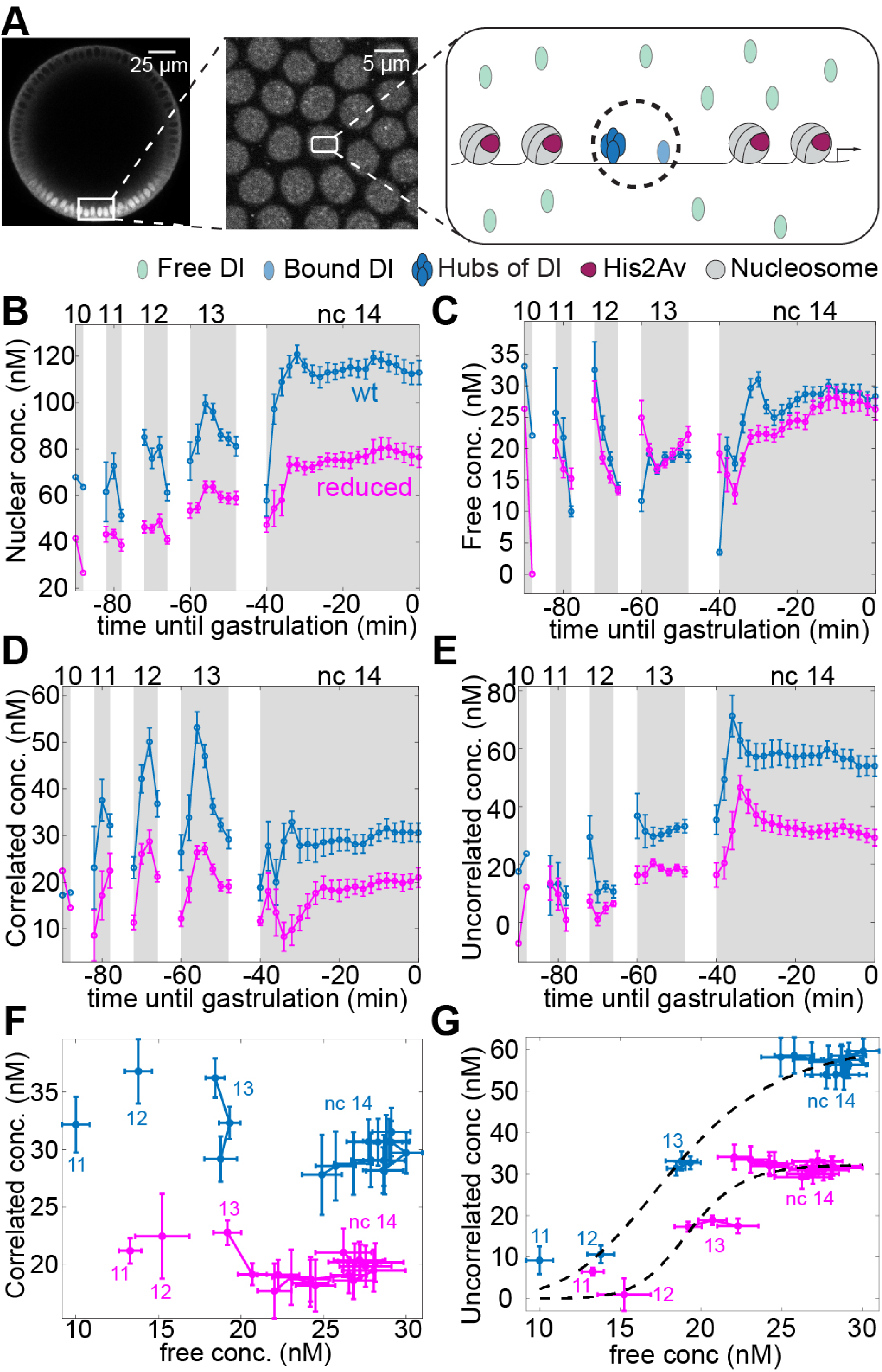
Temporal variation of biophysical parameters of Dl in a fly line having wildtype level of Dl (wt) and in a line where the level is reduced (reduced) in the ventral-most nuclei. (A) Representative image showing the Dl gradient (wt) and RICS acquisition in the ventral-most nuclei. We expect to see three populations: free, DNA-bound, and clusters of Dl as depicted in the figure. Populations inside the dashed circle are identified as similar (correlated in this case) using RICS. (B-E) Dynamics of different pools of Dl from nc 10 until gastrulation including total nuclear concentration (B), free concentration (C), correlated concentration (D), and uncorrelated concentration (E). The dynamics of the free population of Dl from nc 10 until gastrulation show only minor differences between the two fly lines compared to the other pools. Curves: mean values (n = 8 for wt embryos, n = 8 for reduced embryos). Error bars: S.E.M. (F-G) Dose/response map between free and the low-diffusivity populations including correlated population (F) and uncorrelated population (G).

According to our RICS measurements, the total nuclear concentration of Dl can be divided into two pools: freely diffusible Dl and slowly-moving Dl (*26, 29, 35*) (Fig. 1A; see Methods). The concentration of the freely diffusible pool of Dl was found to be maintained between roughly 20 and 30 nM during the blastoderm stage, reaching a steady state of ~30 nM during mid nc 14 (Fig. 1C).

The “slowly-moving” pool of Dl, which we defined as having a zero or nearly-zero diffusivity in our RICS analysis, is likely to be composed of Dl monomers or clusters bound to DNA. Both of these species would have a nearly zero diffusivity, and thus are indistinguishable in this method, being lumped together into this pool (indicated by a dashed circle around them in Fig. 1A). This pool could also, in principle, contain slowly-moving Dl with diffusivities significantly less than 1 μm^2^/s, as diffusivities that low would be difficult to distinguish from zero. Therefore, to dissect the slowly-moving pool further, we performed cross-correlation RICS (ccRICS) by cross-correlating the fluctuations between Dl-mNG and His2Av-RFP, (*26, 29, 35*) (see Methods). We found the population of Dl in the slowly-moving pool can be subdivided into the pool of Dl that cross-correlates with His2Av-RFP (likely representing a DNA-bound population (*36, 37*); referred to as the “correlated” pool henceforth), and the pool that does not (referred to as the “uncorrelated” pool) (*35*). The concentration of Dl in the correlated pool peaks in nc 12 and 13, then decreases in nc 14 to a steady state of roughly 30 nM (blue data in Fig. 1D; see also Fig. S1A). From these measurements alone, it is unclear what constitutes the remainder of the slowly-moving pool (not correlating with His2Av). Given the recent discovery of slowly-moving clusters of Bcd and Cic (*33*), we hypothesize the uncorrelated population we observe possibly includes slowly-diffusing oligomeric clusters of Dl (the presence of which is tested in the following sections). The concentration of this uncorrelated pool, determined from the difference between the concentrations of immobile Dl and correlated Dl, increases over time (blue data in Fig. 1E; see also Fig. S1B).

Both sub-populations of the slowly-moving pool are likely due to Dl binding interactions (either to DNA or to some unidentified structures). Therefore, we asked whether these binding interactions follow typical models of protein binding, such as TF/DNA interactions. These models are often loosely based on equilibrium binding affinity and are typically saturable functions of the free concentration, such as Hill functions (*35, 38–42*). In reality, the relation is complicated by the influence of several factors, including binding affinity, chromatin accessibility, and protein-protein interactions (*1–4*). Our measurements of the different sub-populations of Dl afford us the opportunity to test these models of DNA binding by constructing dose/response relationships between the correlated or uncorrelated concentrations of Dl and the free concentration. Surprisingly, we found that the binding of Dl to DNA (correlated concentration) is independent of the free concentration (blue data in Fig. 1F), rather than an increasing function of free concentration (as would be expected). This independence of the free Dl concentration suggests that a simple binding affinity model fails to predict DNA binding, and thus, transcriptional response. In contrast, the uncorrelated pool (blue data in Fig. 1G) could possibly be explained with a dose/response relationship between the uncorrelated concentration and the free concentration. To illustrate this, we fit a 4th-order Hill function to the data (see black dashed curve in Fig. 1G; see also Methods). We noted that, specifically within nc 14, the uncorrelated concentration does not change with variations in the free concentration. This apparent lack of relationship within nc 14 could result from saturation of the uncorrelated pool.

### Reduced levels of Dl result in an altered dose/response relationship

To further test the apparent dose/response relationship, approximated by a Hill function, we asked whether a reduction in Dl levels would result data points that fall on the same dose/response curve. To address this question, we used a Dl-mNG line that has the same protein sequence but has a 50% reduction in Dl levels (referred to as “reduced”; data not shown). Using this reduced fly line is advantageous because we are able to reduce the nuclear Dl levels across the whole time course (magenta in Fig. 1B) while holding the DV location, and hence the Toll-receptor associated signaling (*43–45*), constant.

As the only difference between the wildtype embryos and the reduced embryos is the total Dl concentration (and not Dl protein sequence or DV location), the base expectation would be a reduction in the concentrations of all pools of Dl. While the two slowly-moving populations follow expectations and are generally decreased in the reduced line (magenta data in Fig. 1D, E), the free concentration in the reduced Dl line is, surprisingly, almost identical to that of the wt line (compare magenta and blue curves in Fig. 1C).

If the uncorrelated concentration is truly a simple function of the free concentration, we would expect the data points for both fly lines to fall on the same dose/response curve (uncorrelated concentration vs free concentration). However, we found the data points of the “reduced” embryos were shifted with respect to those from the wt embryos (magenta data in Fig. 1G) rather than falling on the same dose/response curve. To illustrate this, we fit another 4th-order Hill function to the data from the reduced line (see magenta dashed curve in Fig. 1G). As such, a simple dose/response relationship, such as a Hill function dependent only on free Dl concentration, fails to explain the observed behavior, which implies that both sub-populations of slowly-moving Dl are not primarily functions of the available (free) Dl concentration.

Given that the free concentration is the same between the two lines, even though the total concentration differs, our data suggest that there might exist a critical threshold of total nuclear Dl concentration (~30 nM). Below this threshold, all Dl would be freely diffusing, while any Dl concentration above this threshold becomes allocated to the slowly-moving pools (DNA-bound and/or mobile clusters), keeping the free Dl pool unchanged. Such a threshold could be the result of a highly ultrasensitive response to the total Dl concentration (not the free Dl concentration), calling into question whether models of affinity-based binding are appropriate for Dl. Another explanation of this threshold-based behavior would be an LLPS-driven process.

### Free concentration remains unchanged while other populations are dictated by bulk concentration along the Dl gradient

To test whether the relationships we observed also hold across intermediate levels of Dl, and are not simply artifacts of measuring in two different fly lines, we leveraged the fact that the nuclear concentration of Dl varies with DV location. The natural gradient of Dl concentration, which is most intense at the ventral midline (DV coordinate of 0) and decays to near background levels roughly halfway around the embryo (DV coordinate of 0.5; Fig. 2A) (*14, 19*), allows us to measure the effect of continuous variations in total nuclear Dl concentration within the same fly line. Therefore, we imaged different locations along the DV axis during mid-to-late nc 14, when the concentrations of the pools of Dl are each roughly at steady state (Fig. 1B-E; see Methods). We grouped embryos by their DV coordinate, with at least two embryos per bin (width of bin = 0.05).

**Figure 2:**
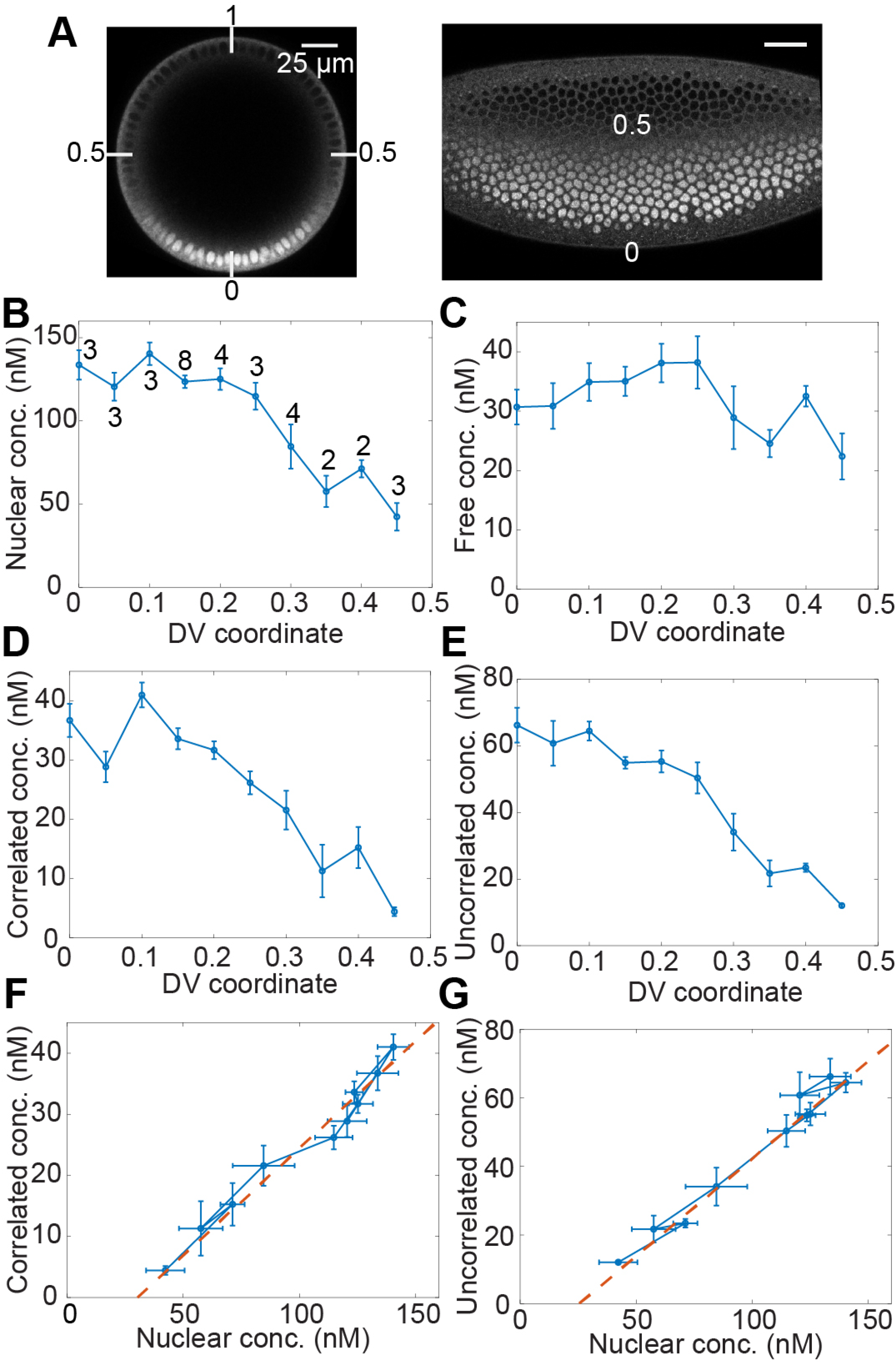
Spatial variation of biophysical parameters of Dl mid nc 14. (A) Snap shots of Dl-mNG in nc 14 embryos, illustrating the relationship between Dl nuclear intensity and DV coordinate in the embryo cross-section (left panel) and surface (right panel). (B-E) Variation of different pools of Dl in ventral to lateral regions of the embryo including total nuclear concentration (B), free concentration (C), correlated concentration (D), and uncorrelated concentration (E). The concentration of free Dl is only a weak function of DV location. Curves: mean values. Error bars: S.E.M. Numbers on curve in (B) indicate sample size at that DV location. (F-G) Dose/response map between total nuclear concentration of Dl and the low-diffusivity populations: correlated population (F) and uncorrelated population (G)

As expected, we found that the total nuclear concentration of Dl decreases as the DV location moves away from the ventral midline (Fig. 2B). The total Dl concentration peaks at the ventral midline (~120 nM) and drops to below 50 nM on the lateral side (DV = 0.45). Interestingly, we noted that the free concentration of Dl has nearly the same value of ~ 30 nM in all the DV locations (Fig. 2C), and that this concentration roughly agrees with the free concentrations we observed for the “wt” and “reduced” lines in the RICS time course (Fig. 1C).

The fraction of both pools of Dl composing the immobile fraction: the correlated fraction representing the DNA-bound pool determined by ccRICS (Fig. S2A) and the uncorrelated fraction representing the pool immobilized likely due to the formation of clusters (Fig. S2B) showed slight reduction with distance form ventral midline.

We found that the concentrations of both slowly-moving pools of Dl decrease with increasing distance from the ventral midline, due to the decrease in the total nuclear concentration of Dl (Fig. 2D and E). This is expected, as a higher concentration of bound Dl is needed on the ventral side for driving the expression of target genes. Taken together with the nearly-constant concentration of free Dl, we found no clear relationship between the concentrations of the slowly-moving pools of Dl and the concentration of free Dl (Fig. S2C,D). In contrast, the relationship between the concentrations of these pools of Dl and the concentration of total Dl is almost linear (Fig. 2F, G). Furthermore, a small extrapolation of these linear relationships reveals an x-intercept of ~30 nM of the total Dl concentration. Thus, our *in vivo* measurements along the DV axis further support that there exists a threshold of total Dl of ~30 nM that must be reached to facilitate the formation of the slowly-moving populations of Dl (either by DNA binding or cluster formation), independent of the DV location. However, we cannot rule out that our observed relationship between total Dl and the slowly-moving pools may be in the linear regime of a highly sigmoidal function.

### Threshold concentrations in Dl possibly indicate thermodynamic phase separation

The presence of a threshold concentration mentioned above is a phenomenon observed in LLPS (*46*). While this is a controversial topic, we asked whether any other hallmarks of LLPS exist in the Dl system. First, we noted that, in a pure-species system that undergoes phase separation, once a threshold concentration is reached, any increase in the total concentration changes relative volumes occupied by the phases, but not their concentrations (*46*). While this is only strictly true of a pure species system, it may be a reasonable approximation for the Dl system if other components do not strongly affect putative phase separation. If so, then the free population (representing the dispersed phase) and the slowly-moving ones (both representing condensed phases) should each have constant concentrations. We observed a constant concentration of the dispersed phase, consistent with LLPS (Figs. 1C, 2C). On the other hand, our measurements of the condensed phase concentrations have been averaged over the whole nuclear volume. Under the assumption of LLPS, these averaged concentrations would be equal to the product of the actual (fixed) concentrations and the volume fraction of the respective condensed phases (see Methods). Furthermore, the volumes occupied by these phases, and thus our measured concentrations, would be directly proportional to the total nuclear concentration of Dl above the threshold of 30 nM, consistent with our observations (Fig. 2F, G; see Methods).

Second, weak multivalent protein-protein interaction mediated by intrinsically disordered regions (IDRs) or low-complexity domains (LCDs) lead to LLPS (*5, 46–49*). We thereby used predictive algorithms such as PSPredictor (*50*), PhaSePred (*51*), FuzDrop (*52*), and PSPHunter (*53*) to estimate the propensity of LLPS in case of Dl. All the algorithms returned a high score indicating that Dl is likely to phase separate (see Table S1), as opposed to mNG, the fluorescent protein tagged to it, which shows very low propensity to phase separate. While the putative threshold and the IDRs intriguingly point to LLPS of Dl, showing that a system in fact phase separates is difficult, and further *in vivo* measurements to infer the material properties of the clusters and the mechanism of formation (*46, 49*) are needed before a definitive case can be made.

### Single particle tracking reveals slowly-moving clusters of Dl

The results so far point to the presence of two pools of Dl with different diffusivities: fast (possibly free Dl) and slow (possibly clusters of Dl and/or DNA-bound Dl, corresponding to the uncorrelated and correlated populations, respectively) (Fig. 1A). Any clusters of Dl, either those known to be bound to DNA (*13, 23*) or the putative clusters in the uncorrelated population, would be brighter than the single molecules of freely disusing Dl. Therefore, we performed single particle tracking using super-resolution imaging to identify and characterize the mobility of the clusters of Dl. We used the Zeiss Airyscan detector that allowed the fast imaging with an improved signal-to-noise ratio (SNR) compared to traditional confocal acquisitions, which is needed for particle tracking experiments (*54*). We imaged the ventral-most nuclei in the embryos at mid nc 14 with a rapid frame rate (90 ms per frame). In the average image over the whole time series, bright puncta were clearly visible for Dl-mNG, whereas no such puncta were observed in control embryos expressing NLS-eGFP (Fig. 3A). The Fiji Mosaic plugin was used to detect and track the particles (*55*) (see Methods), and we found clusters of Dl that lasted for several frames (Fig. 3A and Movie S2). We found that the total number of particles detected during the acquisition time decreased with increasing distance from the ventral nuclei (Fig. 3B). This suggests that more clusters are formed at high levels of Dl. A trend similar to that observed in case of the correlated and uncorrelated populations using RICS (Fig. 2D, E).

**Figure 3:**
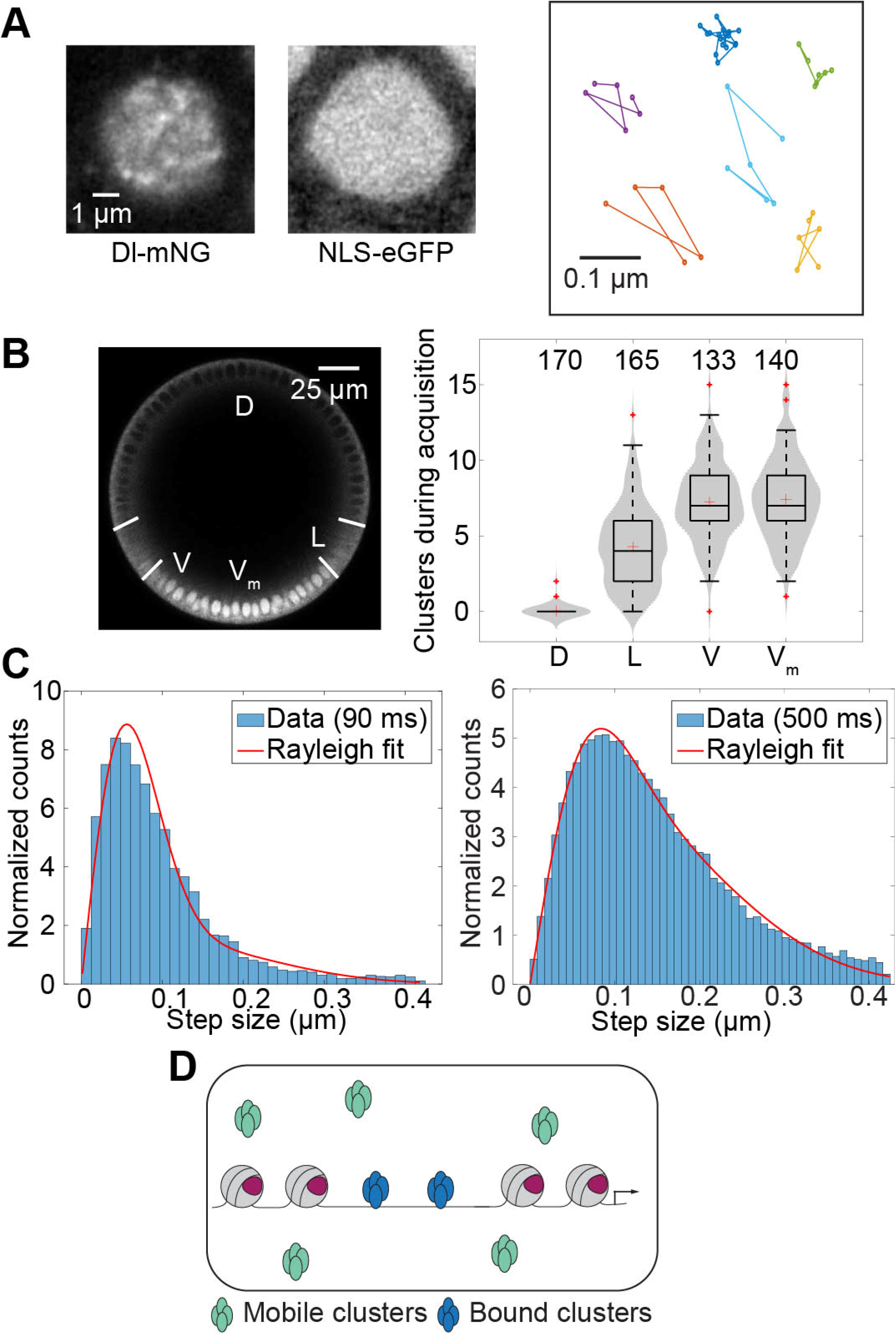
Single particle tracking to characterize Dl clusters. (A) Representative average image of a time series for Dl-mNG and NLS-eGFP and representative trajectories of the particles. (B) Number of clusters detected during acquisition on different DV locations as showed in the left panel. Numbers in the right panel indicate sample size. (C) Step size distribution and 2-component model fitting for the determination of diffusivities and compositions of different populations for trajectories detected in 90 ms frame rate (left panel, n = 140 nuclei) and 500 ms frame rate (right panel, n = 118 nuclei). (D) Bound and slowly-moving clusters are identified using SPT.

To analyze the movement of these particles, we used a step size distribution (SSD) approach, in which the step sizes of particles between two consecutive frames are recorded. The shape of the distribution of these step sizes can be analyzed to determine not only the diffusivities of the particles, but also whether multiple sub-populations with different diffusivities are present (*56*). Following this approach, we fitted a two-component Rayleigh distribution (see Methods) to the distribution of the step sizes (*33*) and found two populations with diffusion coefficients of 0.1 µm^2^/s and 0.01 µm^2^/s (27% and 73% of the total clusters respectively; Fig. 3C). These diffusivities are very similar to those recently reported for DNA-bound hubs and for mobile (slowly-moving) clusters, respectively, for the early *Drosophila* transcription factors Bcd and Cic (*33*). As such, we interpret the slower population with 0.01 µm^2^/s diffusivity to be a DNA-bound population of Dl and the relatively faster population with 0.1 µm^2^/s diffusivity to be slowly-moving clusters of Dl. To determine whether the two-component model is the best fit, we also fitted one- and three-component distribution models to our data. The one-component distribution model could not explain the data (see Fig. S3A), and, in the three-component model, the first two components match those of the two-component model, while the third component is optimized to be zero percent of the total population (Fig. S3A). Notably, the population of free Dl (D ~ 3-4 μm^2^/s (*29*)) is not detected in this experiment because free Dl would not form bright clusters and because the frame time is too long to detect a free diffuser. This suggests that, at a long enough frame time, slowly-moving clusters will be also blurred into the background and only the DNA-bound population will be detected (*10, 57*). Indeed, when we used a longer frame time of 500 ms, we could only detect the population of particles bound to DNA (Fig. 3C).

Mean-squared displacement (MSD) (*58, 59*) analysis, although only applicable for the determination of diffusivity when a single population is expected, can still provide qualitative information about the type of motion of the particles. The slope in the MSD vs. Δt plot and slope of the MSD/Δt vs. Δt plot, where Δt is the lag time, provide insights into the diffusivity regime of the particles. For the 500 ms frame time, we found the slope to be less than 1 in the MSD vs. Δt plot and negative in the MSD/Δt vs. Δt plot (Fig. S3B), indicating the presence of the population in the sub-diffusion regime, which results from confined diffusion, consistent with movement of particles attached to the DNA (*10, 55, 60, 61*). Taken together, these results suggest that there are at least two separate populations of clusters of Dl: DNA-bound and the slowly diffusing (Fig. 3D).

### Modified RICS imaging parameters simultaneously detect three populations

Our RICS experiments described in a previous section showed there is a free population of Dl-mNG and a slowly-moving population (*29*). Using SPT, we found that the slowly-moving population is composed of at least two sub-populations: one bound to the DNA and another of slowly-moving Dl clusters with a diffusivity of roughly D = 0.1 µm^2^/s. As such, the SPT and RICS analyses together have detected a total of at least three different populations on the basis of diffusivity: freely diffusing Dl, slowly-moving clusters, and DNA-bound Dl. However, the individual experiments could not detect all three populations together, as the free population detected in RICS moves too quickly to be detectable with SPT, while the slowly-moving sub-populations cannot be readily distinguished in our previous RICS experiments.

To simultaneously detect all three populations (Fig. 4A), and, in particular, to test the hypothesis that there is a slowly-moving population of Dl, distinguishable from truly immobile Dl, we used a longer line time for the RICS acquisitions. The longer line time would give the slowly-moving clusters more time to diffuse to a new location during the raster scan, thus allowing us to detect them, while still allowing us to detect the free Dl. Therefore, to alter the line time while holding as many other imaging parameters (such as pixel dwell time, image size, and frame time) as constant as possible, we imaged 4096 x 1024 pixel areas (133 μm x 33 μm) along the ventral S3). In Case 1, we oriented the image acquisition vertically, such that the line time was τ ℓ ~ 5 midline of nine nc 14 embryos multiple times under two different Cases (see Methods and Movie ms (Fig. 4B), consistent with the line time in our RICS acquisitions described in a previous section. In Case 2, we flipped the orientation of the image acquisition, resulting in a line time of τ ℓ ~ 17 ms (Fig. 4C), which should be able to detect particles with a diffusivity as low as roughly 0.1 μm^2^/s.

**Figure 4:**
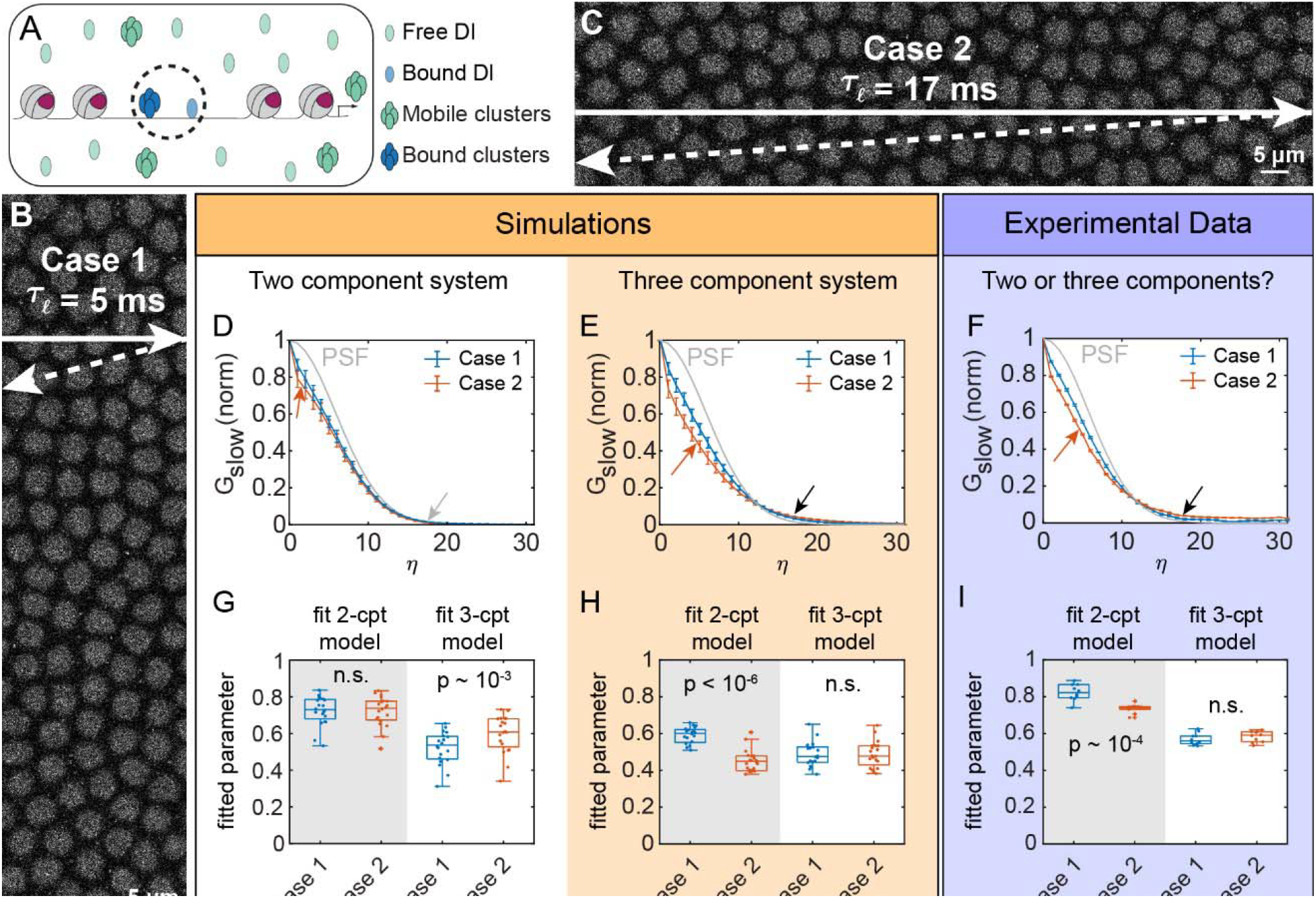
Variation of the line time of the RICS experiments to capture the slowly-moving clusters of Dl. (A) Populations present as indicated by the variation of the line time of the RICS experiments. Populations inside the dashed circle are identified as similar (bound in this case) in this experiment. (B) Case 1: Acquisition of a 133 μm x 33 μm area of the embryo along the ventral midline, oriented such that the line time is roughly 5 ms. Arrows denote exaggerated laser path: the horizontal arrow represents line scanning, and the dashed diagonal arrow represents line retracing. Together, the lengths of these arrows roughly correspond to the line time. (C) Case 2: The same embryo is imaged with the microscope axes reoriented by 90°, ensuring the line time is much longer (17 ms). The arrows are similar to those in (A), illustrating the longer line time in Case 2. (D) Simulations of a two-component system (free Dl + immobile Dl) showing the averaged, normalized slow-direction ACFs for Cases 1 and 2. The simulated PSF is shown in gray. The Case 2 ACF visually differs from Case 1 only slightly and only at pixel shifts 1 and 2. Neither ACF extends past the PSF (gray arrow). Number of simulations for each Case: 20. Error bars: SEM. (E) Simulations of a three-component system (free Dl + slowly-moving clusters + immobile Dl) showing the averaged, normalized slow-direction ACFs for Cases 1 and 2. The simulated PSF is shown in gray. The ACFs of the two Cases clearly differ at intermediate pixel shifts (η between 1 and 10, red arrow), and both ACFs extend past the PSF (black arrow). Number of simulations for each Case: 20. Error bars: SEM. (F) Experimental data with unknown number of components. The weighted average of normalized slow-direction ACFs for Cases 1 and 2 differ significantly at pixel shifts 1-9 (red arrow). Both ACFs have tails that extend past the PSF (black arrow). N = 9 embryos with at least two acquisitions of each Case. Error bars: weighted SEM. (G) Model fits to the simulated two-component system. Fitting a two-component model to a two-component system produced the same values of fitted parameter between Cases 1 and 2. However, mismatching a three-component model fitted to a two-component system produced statistically different values of the fitted parameter between the two Cases. (H) Model fits to the simulated three-component system. Mismatching a two-component model to a two-component system produced statistically different values of the fitted parameter between Cases 1 and 2. However, a three-component model fitted to a three-component system produced statistically indistinguishable values of the fitted parameter between the two Cases. (I) Two- and three-component models fitted to the experimental data. The fits to the two-component model produced statistically different values of the fitted parameter between Cases 1 and 2. However, fitting the three-component model produced statistically indistinguishable values of the fitted parameter between the two Cases.

To determine the expected behavior of the system if slowly-moving clusters are present, we simulated a two-component system (with a freely diffusing population and a truly immobile population, but no clusters), and a three-component system in which slowly-moving clusters are present (see Fig. S4A,B). Our simulations showed that, if slowly-moving clusters are not present, there would be very little difference in the slow direction cut of the ACF between Cases 1 and 2 (Fig. 4D, “Two component system”; see also Methods), only a small deviation at low pixel shifts (see red arrow in Fig. 4D). In addition, the large-pixel-shift tails of the ACF should not extend past the PSF (see gray arrow in Fig. 4D). On the other hand, if there is also a third population of slowly diffusing clusters of Dl, there should be a clear difference in the slow direction cut of the ACF between Cases 1 and 2 (compare blue curve to the red curve in Fig. 4E, “Three component system”). Furthermore, the ACFs of both Cases should extend beyond the tail of the PSF at large pixel shifts (black arrow in Fig. 4E). Our experimental results show that the slow direction of the ACF is clearly different between Cases 1 and 2, and both extend past the PSF, suggesting there are indeed three populations with different diffusivities present (Fig. 4F).

To quantitatively test our hypothesis that there is a slowly-moving population of Dl, we fit two- and three-component models to our data. If we fit these models to the simulations of a two-component system, the fitted parameter of the two-component model (the fraction slowly-moving) would not be statistically distinct between Cases 1 and 2. However, the fitted parameter in the three-component model (the fraction of mobile clusters) would be statistically distinct between Cases 1 and 2 (Fig. 4G), because the three-component model would not match the underlying two-component system. In contrast, if we fit these models to the simulations of a three-component system, the opposite would be true (Fig. 4H). In other words, if the model matches the system, fitted parameters should match between Cases 1 and 2, while if the model is a mismatch to the system, the fitted parameters should also be mismatched between Cases 1 and 2. When we fit the two models to our experimental data, we found a clear statistical difference between Cases 1 and 2 for the two-component model fit, but no statistical difference between Cases 1 and 2 for the three-component model fit (Fig. 4I). Therefore, the combination of RICS analyses for Cases 1 and 2 clearly allow for simultaneous detection of all three populations of Dl, including the slowly-moving clusters, which might be formed due to LLPS (Fig. 4A).

Furthermore, our three-component model fits suggest that these slowly-moving clusters have a weighting *ϕ*_1_ =0.6 These results further verify our observations for the spatiotemporal variation using RICS and the SPT results.

## Discussion

Despite decades of research, the molecular mechanisms behind the precise gene regulation by sequence-specific transcription factors, particularly mechanisms linking nuclear concentration of the transcription factor to its DNA binding (a crucial step in gene regulation), remain unresolved (*22, 62*). Transcriptional regulation is dependent on TF dynamics such as the total concentration, fraction bound, and hub formation (*5*). Therefore, in this study, we investigated how the bulk concentration of Dl relates to DNA binding and hub formation. Using experiments that span multiple length and time scales, we detected at least three different populations of Dl having distinct diffusivities present in the nuclei: freely diffusing (D ~ 3 µm^2^), previously unobserved slowly-moving clusters (D ~ 0.1 µm^2^), and a DNA-bound population (D ~ 0.01 µm^2^; see Fig. 5).

**Figure 5:**
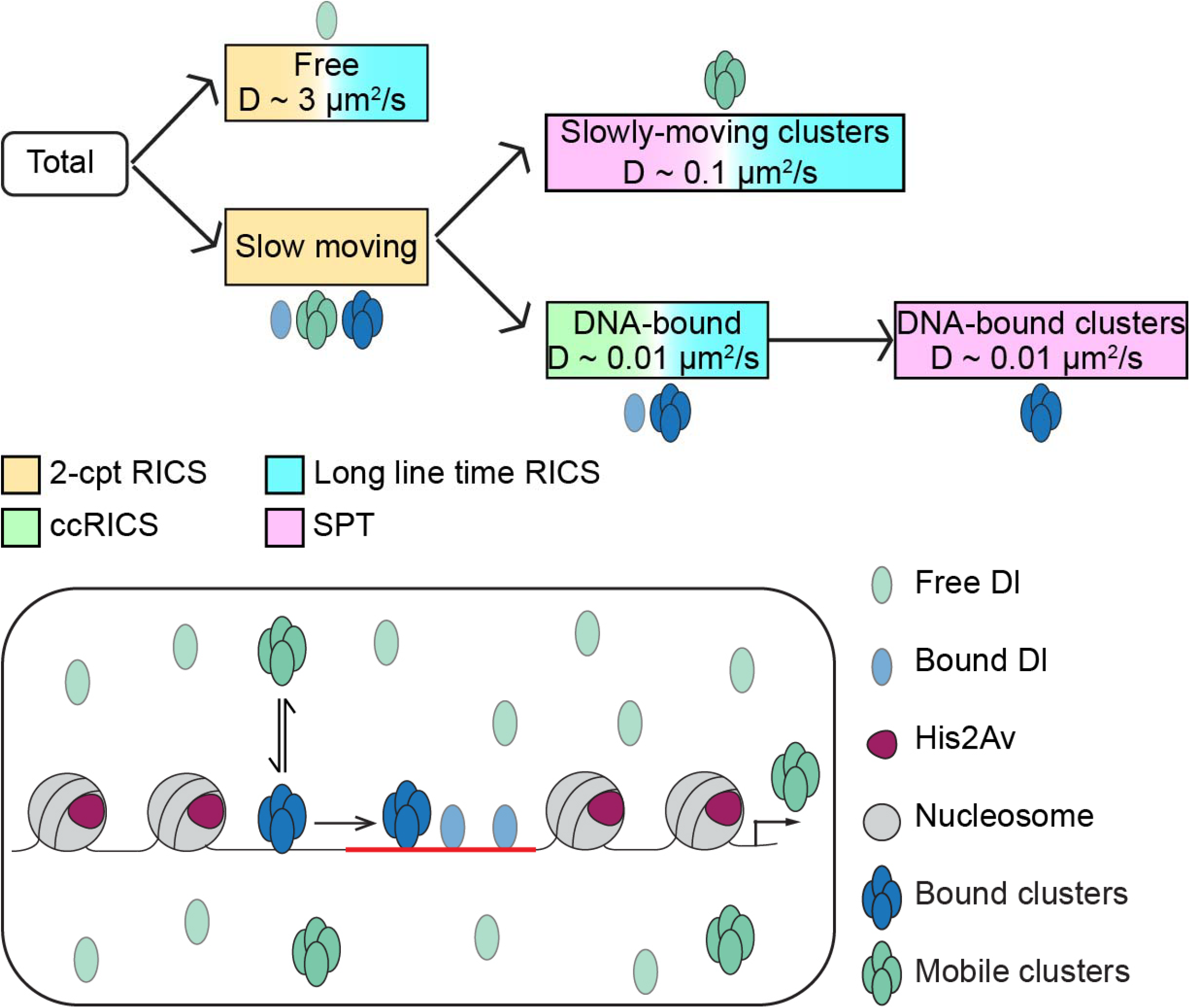
Model of Dl-DNA binding as indicated by multiple experimental techniques. The total population of Dl in the nuclei is composed of free and slow populations as indicated by RICS. The slow population is composed of DNA-bound population (indicated by ccRICS) and slowly-moving clusters. SPT can differentiate between the DNA-bound clusters and slowly-moving clusters. Finally, long line time RICS can detect the presence of all the constituent populations together. Both the experimental techniques and the different populations are color coded.

Recent work has suggested that clusters, or hubs, of transcription factors are significant in DNA binding. For example, DNA-bound hubs of the morphogen Bcd have been proposed recently to facilitate the transcriptional activity by locally increasing the concentration (*10, 11*) or by acting as fast sensor of concentration (*63*). Similarly, enrichment of puncta of Dl near individual transcription sites has been observed (*13, 23*). A recent study has shown that Bcd and Capicua (Cic), two TFs active in the early *Drosophila* embryo, when Dl is also active, form slowly-moving aggregates (D ~ 0.1 μm^2^/s), which might provide an efficient search mechanism for the target sites (*33*). In this work, we found a previously-unobserved population of slowly-moving clusters of Dl, suggesting that this might be a common mechanism of transcriptional regulation by morphogen gradients. Thus, gene regulation models in eukaryotes must take the formation of DNA-bound as well as slowly-moving hubs of TFs into account.

In addition to identifying three sub-populations of Dl, we also leveraged the variation of Dl concentration across nc 11-14, along the DV axis, and in two different fly lines to map the relationships among them. Strikingly, we found the free concentration of Dl remained roughly the same (~30 nM), regardless of the bulk concentration, indicating the possibility of a threshold concentration of total Dl, above which further increases in Dl became exclusively allocated to the slowly-moving populations. Further supporting the presence of a threshold in bulk concentration, the dose/response maps connecting the two slowly-moving populations to the total Dl concentration were linear, extrapolating back to this same 30 nM threshold in total Dl concentration. Thus, our nucleus-level measurements, while not as specific as those that focus on individual enhancers (*13*), have provided crucial details regarding the mechanisms of DNA binding and cluster formation.

The presence of a threshold in the total nuclear concentration of Dl, above which no further free Dl is allocated, is a hallmark LLPS of biomolecular condensates (*64*). While the clusters of Dl forming from LLPS is an intriguing possibility, other work suggests the opposite, and overall, LLPS of transcription factors has been a hotly-debated topic in the past decade (*48, 65–67*). LLPS has been proposed to play a role in buffering biological fluctuations, aiding in robustness of certain pathways, or repressing pathways whose functioning requires noise (*48, 68*). However, quantitative methods to provide definitive evidence of LLPS are still lacking in the field (reviewed in (*49*)). Therefore, regardless of the mechanistic status of Dl cluster formation, our *in vivo* quantitative measurements and methodologies provide a general, quantitative framework for testing properties of clusters for LLPS-like behavior by the detecting relationships among the constituent populations *in vivo*.

While may subpopulations detected in one experiment are similar to those detected in other experiments, we acknowledge these similar subpopulations may not necessarily be identical across experiments. For example, the correlated and uncorrelated populations from RICS/ccRICS experiments may not exactly map one-to-one onto the DNA-bound clusters (0.01 μm^2^/s) and slowly-moving clusters (0.1 μm^2^/s) detected in the SPT experiments, as some of the correlated RICS population could be DNA-bound monomers or dimers of Dl. Additionally, we acknowledge the possibility that there may be populations having intermediate composition and diffusivity not identified in our experiments. One further caveat is that clusters of fluorescently-labeled molecules, by definition, have a higher molecular brightness than the monomers and dimers, which affect the quantitative values of the RICS analyses(*27, 35, 69*). In particular, corrections for differing brightness values would result in a slight increase to the total concentration of Dl, and the concentration of the other subpopulations would also scale accordingly. Even so, our general findings, including the discovery of mobile clusters of Dl and the linear/threshold relationship of slowly-moving populations to the total Dl concentration, are not affected by these caveats.

The existence of clusters, both DNA-bound and mobile, have profound implications for our understanding of gene regulation. Put within the broader context, these clusters must be accounted for along with other crucial parameters that explain binding of a TF, such as the chromatin accessibility and cooperativity with pioneer-like factors. All together, our results support a model in which, during the early ncs, owing to its lower concentration, Dl binds only the accessible sites created by pioneer factors. During nc 14, when the nuclear concentration of Dl is high, binding is driven by the concentration of Dl, perhaps by clusters of Dl. This is in agreement with the proposed cooperativity between the pioneer factor Zld and Dl and the sporadic/delayed expression of Dl target genes in Zld mutant embryos (*40, 70–72*). Our results also suggest that, during nc 14, binding occurs when a threshold concentration of free Dl is exceeded. As such, different populations of Dl at varying concentrations within and across nuclei must be taken into account to understand its concentration-dependent transcriptional interpretation. More broadly, we anticipate that similar detailed measurements of nuclear subpopulations will be needed for other transcription factor systems, potentially pointing to generalizable rules of transcriptional regulation (*10, 11, 13, 33, 35, 63, 73*).

## Materials and Methods

### *Drosophila* strains

The fly strain used for all the imaging for Dl is yw/w; Dorsal-mNeonGreen /cyo; +. For the reduced level of Dl we used, yw/w; Dorsal-mNeonGreen-dsRed/cyo; +. The other fly strains used are: w[*]; P{w[+mC]=His2Av-mRFP1}II.2 (Bloomington Drosophila Stock Center (BDSC) #23651) for nuclear segmentation and ccRICS using His2Av-RFP, w[*]; P{w[+mC]=His2Av-EGFP.C}2/SM6a (BDSC #24163) for using His2Av-eGFP as photobleaching control in the particle tracking experiments.

### *Drosophila* husbandry and sample preparation

The fly stocks were grown on standard cornmeal-molasses-yeast medium at 25°C. Fly cages were set with males and females of desired fly lines two days before imaging and kept at room temperature. Grape juice plates were streaked with yeast paste and placed on the bottoms of the cages for egg laying. The plates were changed once every day. On the day of imaging, the grape juice plates were placed for oviposition for an hour after which it was removed to collect the embryos. The embryos were washed from the plate into a mesh basket using deionized water. They were then dechorionated in bleach for 30 s and washed again with deionized water to remove residual bleach. The embryos were mounted in 1% solution of low melting point agarose (IBI Scientific, IB70051) in deionized water on a glass bottom Petri dish (MatTek, P35G-1.5-20-C). The mounting was done in such a way that the ventral surface of the embryo touched the coverslip in the Petri dish for imaging the ventral nuclei. For imaging other DV positions, the angle at which the embryo touched the coverslip was changed by rotating it slightly with a hair loop. The agarose was let to solidify after which deionized water was poured in the dish to avoid drying of the agarose covering the samples.

## Confocal imaging

### Spatial and temporal variation RICS

All the imaging was performed using LSM 900 (Carl Zeiss, Germany) confocal laser scanning microscope. For the confocal imaging C-Apochromat 40x/1.2 water immersion Korr objective, 488 nm laser for the mNeonGreen, 561 nm laser for RFP, and GaAsP-PMT detector were used. Embryo stage was estimated based on the His2Av-RFP signal. Image acquisition was started when the embryos reached the desired stage: nc 10 for the temporal variation study and mid nc 14 for the spatial variation study. A frame size of 1024 × 1024 pixels at 5x zoom was used. This corresponded to a pixel size of 31.95 nm and a pixel dwell time of 2.06 μs at a frame time of 5.06 s. The acquisition was continued until the embryo gastrulated as indicated by the nuclear morphology.

To minimize photobleaching, the ROI was moved to a different part of the embryo after every 12 frames ensuring that all the locations are aligned in the same DV coordinate. RICS analysis on the 12 frame acquisitions of time series enabled the precise determination of the temporal dynamics of the total nuclear concentration and binding of Dl.

### RICS analysis and fitting of ACF models

We performed RICS analysis according to previously-published protocols (*29, 35*). In brief, two dimensional (2D) RICS autocorrelation functions of the nuclear fraction of each frame were built using a fast Fourier transform protocol in Matlab, and these ACFs were averaged together for all frames in a given group. The result was a time series of 2D ACFs, each of which corresponded to a given grouping of 7-12 frames. Background subtraction was performed on the fly by examining the histogram of intensities and fitting a Gaussian to the lowest intensity pixels in the image.

The theoretical 2D RICS ACF, *G*(Δ*x*, Δ*y*), is the following:

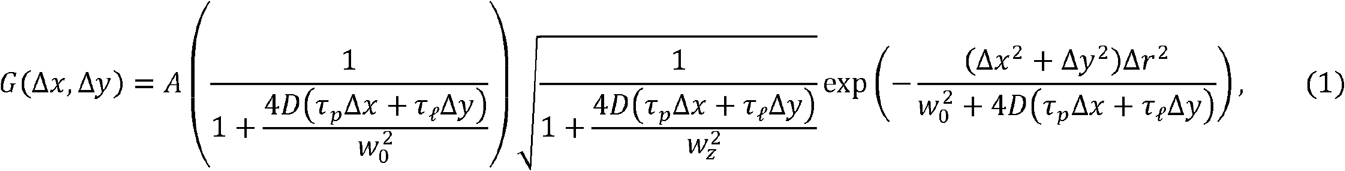

Where *A* is the amplitude; *D* is the diffusivity; *w*_0_ and *w*_*z*_ are the radii of the point spread function (PSF) in the *xy* plane and the axial (*z*) direction, respectively; Δ_*r*_ is the *xy* size of a pixel; τ_*p*_ and τ_*ℓ*_ are the pixel dwell time (determined by the scan speed) and line time (determined by a combination of the scan speed and number of pixels in the width of the image), respectively; and Δ*x* and Δ*y* are the pixel shifts in the fast and slow directions, respectively.

In practice, 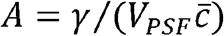, where 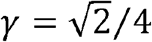 is a factor that accounts for the uneven illumination airy unit, 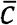 is the average concentration in the confocal volume, and 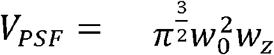 is the effective volume of the point spread function.

The fast direction of the data-derived 2D ACF, at each time point, was used to fit the fast direction cut of the theoretical 2D ACF, *G*(Δ*x*, 0), which reduces to a Gaussian equation that approximates the microscope’s PSF:

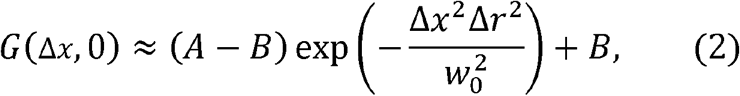

To improve the robustness of the fit, a small, adjustable background constant,*B*, was added and *w*_0_ was allowed to vary slightly. The parameter *B* was constrained to have a magnitude less than 10^-3^. This first fitting step gave us an estimate of the ACF amplitude, *A*, and hence, the average total concentration.

After obtaining an estimate of *A*, we held it fixed and used the entire 2D ACF to fit *G*_2*c*_(Δ*x*, Δ*y*) which is a linear combination of two components, a freely diffusing population and a slowly-moving population (*29*):

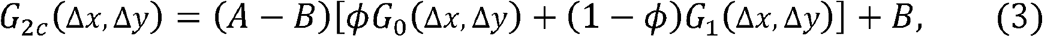

where *G*_0_(Δ*x*, Δ*y*) is given by Equation 1 with *A* = 1 and *D* = 0 *G*_1_(Δ*x*, Δ*y*) is given by Equation 1 with *A* = 1 and *D* non-zero, and the linear combination weight, *ϕ*, is the fraction slowly-moving. Note that the parameter *B* in Equation 3 may have a different value from the one found in Equation 2. To avoid overfitting, we assumed *D* = 3 μm^2^/s.

### Fitting ccRICS cross-correlation functions (CCFs) to estimate correlated binding

We performed analysis of ccRICS according to previously published work (*29, 35*). In brief, the 2D cross correlation function (CCF) between Dl-mNG and His2Av-RFP was computed through a fast Fourier transform protocol in Matlab. This 2D CCF was then used to fit a zero-diffusion 2D model of cross correlation (*29*):

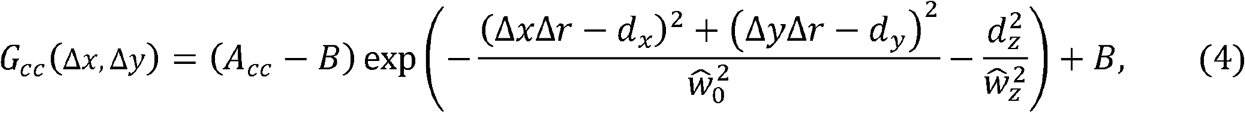

Where *d*_*x*_, *d*_*y*_, *d*_*z*_ represent the displacements between the centers of the two PSFs in the *x, y*, and *z* directions, respectively; 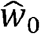 and 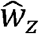 are the average PSF sizes for the planar (xy) and axial (z) directions, respectively; *A*_*cc*_ is the amplitude of the CCF; Δ_*r*_ is the pixel size (see Equation 1); and *B* is a background parameter (see Equations 2 and 3). The axial displacement was previously estimated by imaging fluorescent beads (*35*) and the factor 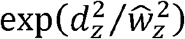 was found to be roughly equal to 1.02, implying the factor could be safely ignored. The average PSF sizes are defined as 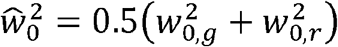 and 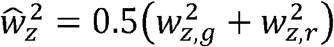, and the subscripts *g* and *r* denote the green and red channels, respectively.

The fraction of Dl-mNG correlated to His2Av-RFP (the correlated fraction) is related to the ratio of *A*_*cc*_ and the ACF amplitude of the red channel, *A*_*red*_:

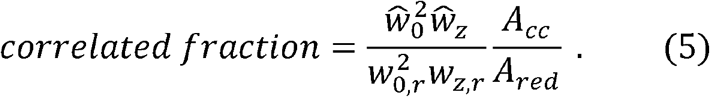

### Relationship between total concentration and condensed phase

Under the assumption of LLPS, to derive the relationship between free Dl (putative dispersed phase) and the correlated and uncorrelated pools of the slowly-moving population (putative condensed phases 1 and 2, respectively), we used the pure species behavior as the foundation. In this case, all phases in equilibrium would have a fixed concentration with respect to varying total concentration, and variations in total concentration would change the volume partitioning among the two phases. We define 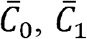, and 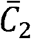 as the fixed concentrations of the dispersed phase and the two condensed phases, respectively. We further define *V*_0,_ *V*_1_ and *V*_2_ as the volumes occupied by the dispersed phase and the two condensed phases, respectively, and note that the total volume of the nucleoplasm available to Dl is *V*_*tot*_ *= V*_0_ + *V*_1_+ *V*_2_.

As our measurements of the free, correlated, and uncorrelated concentrations (*C*_*meas*,O_, *C*_*meas*,1_, and *C*_*meas*,2_, respectively) are averages over the entire volume, *V*_*tot*_, they relate to their respective fixed concentrations by:

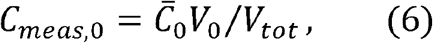

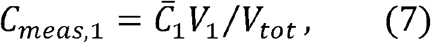

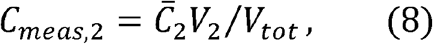

respectively. If *V*_1_+ *V*_2_ ≪ *V*_0_ then Equation (6) reduces to 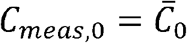, which explains how our measurements of free Dl remain roughly fixed (at 30 nM) with varying total Dl concentration.

The total concentration relates to the phase volumes by:

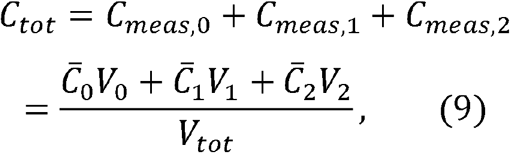

where, in the second line, we have used Equations 6-8. Under the continued assumption that *V*_1_+ *V*_2_ ≪ *V*_0_, Equation 9 can be reduced to:

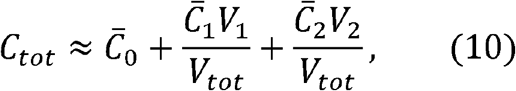

As the three phase concentrations and *V*_*tot*_ are constant, Equation 10 implies that *V*_1_and *V*_2_, and thus also *C*_*meas*,1_and *C*_*meas*,2_ are in a direct linear relationship with the difference between the total concentration, *C*_*tot*_ and the concentration threshold, 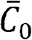.

### Varying line time experiments

In the experiments with varied line times, the pixel size and pixel dwell time were kept the same as the previous experiments at 31.95 nm and 2.06 μs respectively. The line time was varied by changing the frame size. For the shorter line time (Case 1), the frame size was 1024 × 4096 pixels, resulting in a line time of 5 ms. For the same embryo, the longer line time (Case 2) was obtained by rotating the ROI by 90° (exchanging the x- and y-directions of the image), resulting in a frame size of 4096 × 1024 pixels and a line time of 17 ms. This ensured that the imaging parameters will be conserved and each embryo will be imaged with two different line times, the short and the long one, at the exact same DV position. Thus, with the same number of pixels, pixel size, and pixel dwell time the images with frame size of 4096 x 1024 pixels have a line time just under 4 times longer than that of the images with 1024 x 4096 for the same DV location. To minimize and account for photobleaching, the ROI was rotated after 12 frames and for some of the embryos the longer line time experiments were performed first and then the shorter line time ones whereas for the other embryos the order was reversed.

### Simulations of varying line time experiments

We simulated the ACFs of both Cases (the short line time and long line time) for two different systems. The first system was a two component system with freely diffusing Dl-mNG and DNA-bound Dl-mNG (zero diffusion), simulated using Equation 3. We set the baseline diffusivity of free Dl to be 3 μm^2^/s and the baseline slowly-moving fraction (*ϕ*) to be 0.75.

The second system was a three component system with the extra component being a slowly-moving cluster of Dl-mNG. In this system, we assumed the slowly-moving fraction, set at 0.75 for the two-component system, was split into a baseline of *ϕ*_0_ = 0.25 for the fraction bound to DNA and a baseline of *ϕ*_1_ = 0.50 for the fraction of slowly-moving clusters, consistent with our experimental measurements of the correlated and uncorrelated fractions (see Fig. S1):

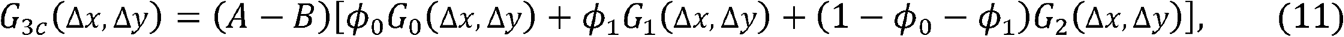

where *G*_0_, *G*_1_, *G*_2_ are each given by Equation 1 with *A* = 1 and with baseline diffusivities of zero, 0.1 μm^2^/s, and 3 μm^2^/s, respectively.

We ran 20 simulations of each system with 10% random noise added to the parameters. After the simulated ACFs were constructed according to Equation 3 (two-component system) or Equation 11 (three-component system), we added 2% Gaussian noise to the ACFs simulate experimental error. The smoothness of the resulting ACFs were comparable by-eye to the smoothness of our typical experimental ACFs. The means of the 20 simulations are found in Fig. 4C and D, with error bars being the standard deviation.

We fit each of these simulations to either a two-component model (Equation 3) or a three-component model (Equation 11). To avoid overfitting, we allowed only one fitted parameter in each case: *ϕ* for the two-component model and *ϕ*_1_ for the three-component model. In both cases, we fixed the diffusivity of free Dl-mNG to be 3 μm^2^/s (*29*). For the three-component model, we additionally fixed *ϕ*_0_ to be 0.25 and the diffusivity of slowly-moving clusters to be 0.1 μm^2^ /s.

### RICS analysis of varying line time experiments and fitting of ACF models

The RICS analysis of our time course data for Cases 1 and 2 in the varying line time experiments was performed as described above. Once the data-derived ACFs were computed, they were used to fit a two-component model (Equation 3) and a three-component model (Equation 11). As described above, to avoid overfitting, we held fixed all parameters except for *ϕ* for the two-component model and *ϕ*_l_ for the three-component model, and we fixed the diffusivity of free Dl-mNG to be 3 μm^2^/s and the diffusivity of slowly-moving clusters to be 0.1 μm^2^/s. We fixed *ϕ*_0_ to be equal to the value of correlated fraction for that time course.

### Super resolution imaging

The particle tracking experiments require rapid frame rates and high spatial resolution, and as such, we used the fast Airyscan detector for improved spatial resolution and signal-to-noise ratio (SNR) (*54*). All the images were taken on the ventral side of the embryos at mid nc 14. Plan-Apochromat 63x/1.4 Oil DIC M27 objective and 488 nm excitation laser for the Dl-mNeonGreen were used. All the images have a pixel size of 42 nm. For the step size distribution experiments the rapid frame rate of 87 ms was obtained by having a frame size of 167 × 167 pixels and the timeseries consisted of 35 frames. Whereas the longer duration of 500 ms was obtained by having a frame size of 380 × 380 pixels and the timeseries consisted of 50 frames to detect the DNA-bound clusters that last for a longer time.

### Particle detection and tracking

The Fiji Mosaic plugin was used for particle detection and tracking (*55*), using the following parameters: particle radius 6 pixels, cutoff score 0%, intensity percentile 0.22%. For particle linking, the maximum step length was set at 10 pixels and the link range 1 frame. Any trajectory lasting less than three steps was discarded.

### Localization precision

To determine the localization precision, we imaged immobilized fluorescent beads (*33, 74, 75*). The beads (Fluoresbrite® YO Carboxylate Microspheres 0.20 µm, Polysciences, Inc., Catalog No. 19391-10) were washed twice with deionized water and then mounted in low melting point agarose on a glass bottom Petri dish in the same way as the embryos to emulate the experimental conditions. The imaging was then performed using the same microscope parameters as those used for the embryos for the 87 ms frame rate. Since localization precision depends on SNR (*76, 77*) which in turn depends on laser power, we used 8 different laser power levels for the imaging: 0.025%, 0.05%, 0.075%, 0.1%, 0.2%, 0.3%, 0.5%, and 0.8%. For each laser power, 10 different time series having 35 frames each at 10 different locations were acquired. The mean diffusivity was calculated using the mean squared displacement (MSD) method (*33, 74*) instead of the step size distribution method since the beads are immobilized thus expected to have long-term trajectories and we do not expect multiple populations with different diffusivities. We calculated the localization precision for all the acquisitions from the diffusivity (*σ*^2^ =4*D*Δ*T*) obtained from the Fiji Mosaic plugin (*33, 55, 74*). The localization precision as a function of SNR was then empirically fit to the following equation (*55, 77*):

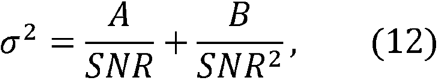

where the first term is due to background noise and the second term is due to shot noise. SNR is defined as the ratio of the mean intensity of the signal and standard deviation of the background. The constants A and B were determined by fitting Equation 12 to the data in the plot of σ^2^ vs SNR plot using lsqcurvefit solver in Matlab (Fig. S3C). The mean intensity of the signal in the embryos was determined at the coordinates of detected particles obtained from the Mosaic plugin and the standard deviation of the intensity outside the detected particles inside the nucleus was determined. The ratio gives the SNR for our samples. The localization precision was then calculated using Equation 12 at the determined SNR.

### Step size distribution

The possibility that multiple populations with different diffusivities might be present and the presence of trajectories exhibiting short-term motion led us to use the step size distribution for the estimation of diffusivities. The length of the steps taken between consecutive frames were calculated and the step size distribution of the particles was obtained. The probability density function of the step sizes was fit with a two-component Rayleigh mixture model (*10, 33, 58, 78*):

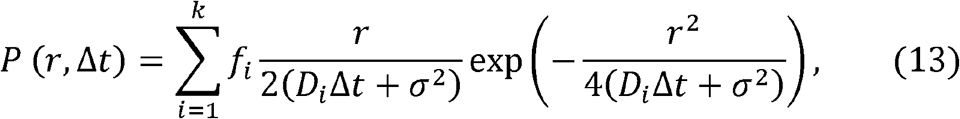

Where *f*_*i*_ is the fraction of the population i (and thereby 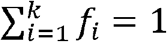), *D_i_* is the diffusivity of the population *i*, Δ*t* is the time between the steps, the localization precision (from the immobilized fluorescent beads experiments) is *σ*^2^ = 4.5 × 10^−4^ and r is the step size. The number of components was changed to 1 and 3 (Fig. S3A) to justify the use of two-component model. The values of diffusivities and fraction of the population in Equation 13 were estimated using the Matlab optimization function fmincon.

## Acknowledgments

S.S.D. was supported by NIH R01 Award R01GM151409 and by NSF Award MCB-2105619. M.A.T. was supported by the National Science Foundation’s Graduate Research Fellowship Program (NSF GRFP).

H.G.G. was supported by NIH R01 Awards R01GM139913 and R01GM152815, by the Koret-UC Berkeley-Tel Aviv University Initiative in Computational Biology and Bioinformatics, by a Winkler Scholar Faculty Award, and by the Chan Zuckerberg Initiative Grant CZIF2024-010479. H.G.G. is also a Chan Zuckerberg Biohub Investigator (Biohub – San Francisco).

G.T.R. was supported by NIH R01 Award R01GM151409 and by NSF Award MCB-2105619.

## Author contributions

Conceptualization: S.S.D and G.T.R Imaging and analysis: S.S.D and G.T.R Generation of fly lines: All authors Investigation: S.S.D and G.T.R Visualization: S.S.D and G.T.R Supervision: G.T.R Writing—original draft: S.S.D and G.T.R Writing—review & editing: All authors

## Data and materials availability

Image analysis pipeline: https://github.com/gtreeves/RICS_timecourse_pipeline All the image files (.czi) will be deposited in the “Texas Data Repository”.

## Supplementary Materials

### Figures

**Figure S1:** Dynamics of fraction of different pools of Dl from nc 10 until gastrulation for wt and reduced fly lines.

**Figure S2:** Variation of fraction of different pools of Dl mid nc 14 at different DV locations.

**Figure S3:** Model fits to the particle tracking parameters.

**Figure S4:** Simulations of individual ACFs.

### Tables

**Table S1:** Phase separation propensity scores of Dl and mNG, the fluorescent protein tagged to it, using multiple algorithms.

### Movie Files

**Movie S1:** Image acquisition on the ventral side of the embryo from nc 10 until late nc 14 for RICS analysis. Mitosis not shown, as only interphase was used for the analysis.

**Movie S2:** Detection of clusters that stay for multiple frames and the trajectories followed by them.

**Movie S3:** Image acquisitions showing Case 1 (short line time) and Case 2 (long line time) for the simultaneous detection of all three populations using RICS.

