## Supplementary Materials for "Global maps of transcription factor properties reveal threshold-based formation of DNA-bound and mobile clusters"

**Figures**

**Figure S1:**

Dynamics of fraction of different pools of Dl from nc 10 until gastrulation for wt and reduced fly lines.

**Figure S2:** Variation of fraction of different pools of Dl mid nc 14 at different DV locations.

**Figure S3:**

Model fits to the particle tracking parameters.

**Figure S4:**

Simulations of individual ACFs.

**Tables**

**Table S1:** Phase separation propensity scores of Dl and mNG, the fluorescent protein tagged to it, using multiple algorithms.

**Movie Files**

**Movie S1:** Image acquisition on the ventral side of the embryo from nc 10 until late nc 14 for RICS analysis. Mitosis not shown, as only interphase was used for the analysis.

**Movie S2:** Detection of clusters that stay for multiple frames and the trajectories followed by them.

**Movie S3:** Image acquisitions showing Case 1 (short line time) and Case 2 (long line time) for the simultaneous detection of all three populations using RICS.

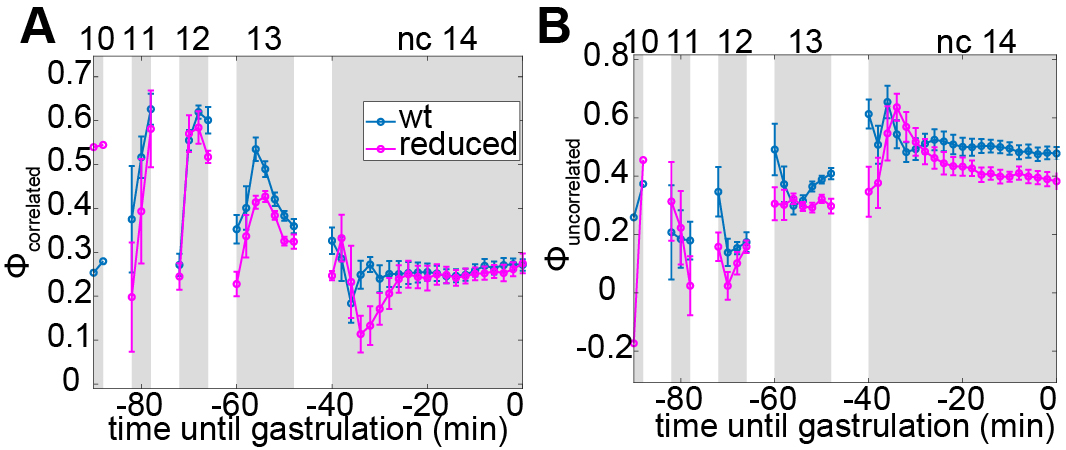

**Figure S1:**

Dynamics of fraction of different pools of Dl from nc 10 until gastrulation for wt and reduced fly lines.

(A) Fraction of correlated population
(B) Fraction of uncorrelated population

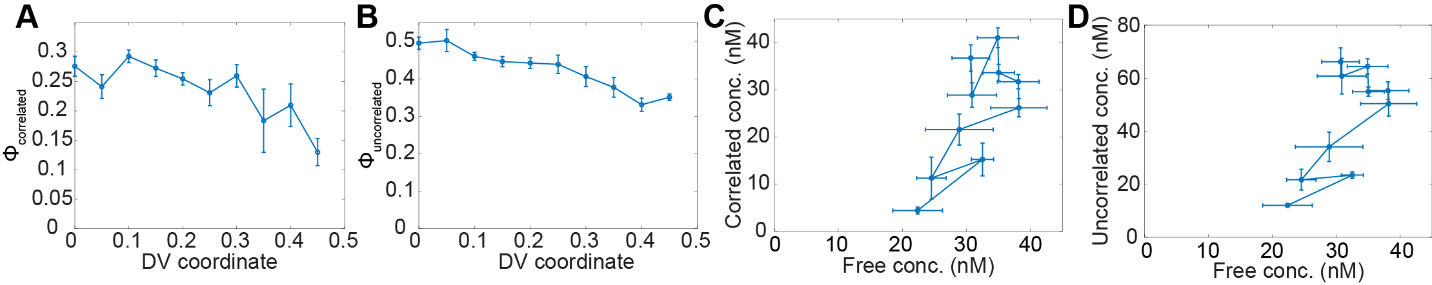

**Figure S2:** Variation of fraction of different pools of Dl mid nc 14 at different DV locations.

(A) Fraction of correlated population
(B) Fraction of uncorrelated population

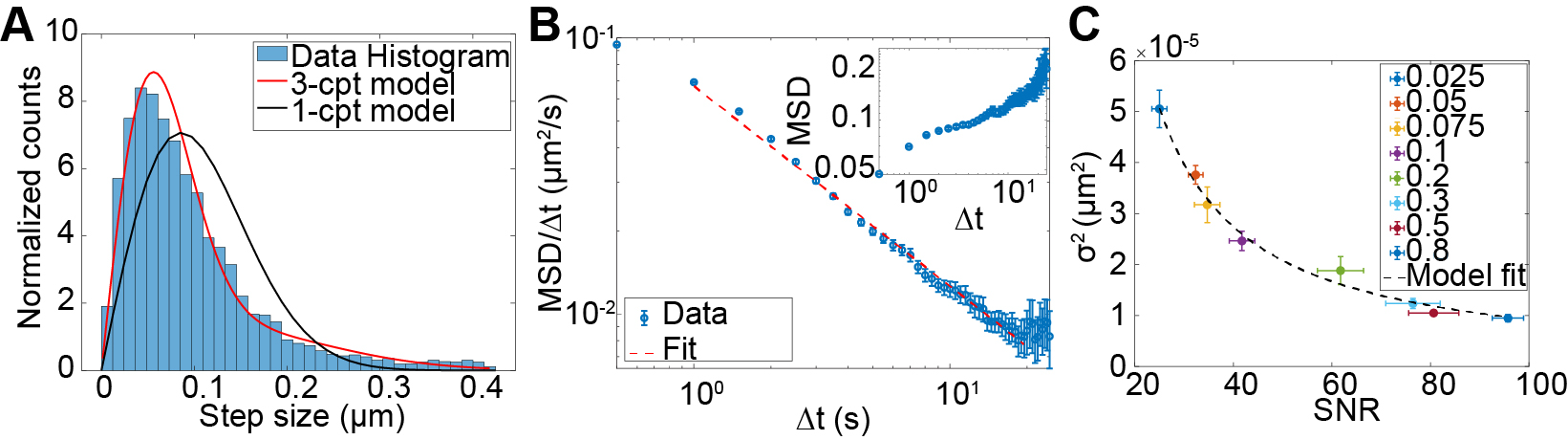

**Figure S3:**

Model fits to the particle tracking parameters.

(A) 1- and 3-component model fitting for the determination of diffusivities and compositions of different populations
(B) MSD analysis for the qualitative estimation of the diffusion regime of the population detected in the longer frame rate of 500 ms
(C) Determination of localization precision using immobilized fluorescent beads imaged with varying laser powers. Each laser power was used to obtain 10 different acquisitions

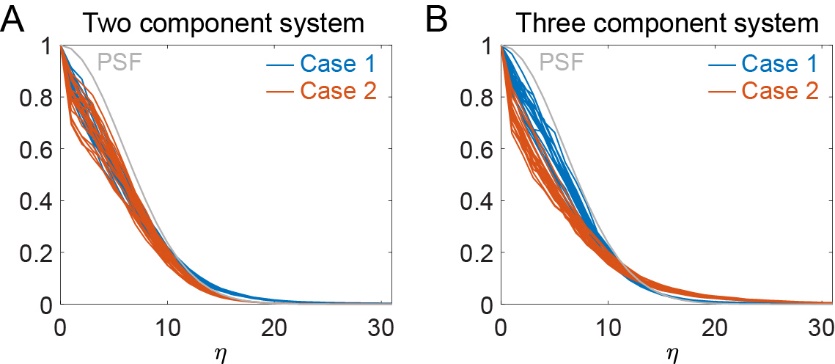

**Figure S4:**

Simulations of individual ACFs.

(A) Twenty simulations of a two-component system for Case 1 (short line time; blue curves) and Case 2 (long line time; red curves).
(B) Twenty simulations of a three-component system for Case 1 (short line time; blue curves) and Case 2 (long line time; red curves).

**Table S1:** Phase separation propensity scores of Dl and mNG, the fluorescent protein tagged to it, using multiple algorithms.

| Algorithm | Score for Dl | Score for mNG | Description |
| --- | --- | --- | --- |
| PSPredictor (*50*) | 0.99 | 0.0815 | Phase separation protein prediction score, highest being 1 and the lowest 0 |
| PhaSePred (*51*) | 0.93 | - | Rank score, where highest rank score is 1 and the lowest is 0 |
| FuzDrop (*52*) | 0.99 | 0.2291 | Probability of spontaneous liquid-liquid phase separation |
| PSPHunter (*53*) | 0.81 | 0.55974 | Probability to undergo phase separation |
